# Effects of TMS over the anterior intraparietal area on anticipatory fingertip force scaling and the size-weight illusion

**DOI:** 10.1101/2020.05.18.101675

**Authors:** Vonne van Polanen, Gavin Buckingham, Marco Davare

## Abstract

In skilled object lifting, fingertip forces need to be carefully scaled to object weight, which can be inferred from object properties, such as size or material. This anticipatory force scaling ensures smooth and efficient lifting movements. However, even with accurate motor plans, weight perception can still be biased. In the size-weight illusion, objects of different size but equal weight are perceived to differ in heaviness, with the small object perceived to be heavier than the large object. The neural underpinnings of the size-weight illusion and anticipatory force scaling to object size are largely unknown. In this study, we hypothesized a possible role of the anterior intraparietal cortex (aIPS) in predictive force scaling and the size-weight illusion, which we investigated by applying continuous theta burst stimulation (cTBS) prior to participants lifting objects designed to induce the size-weight illusion. Participants received cTBS over aIPS, the primary motor cortex (control area), or sham stimulation. We found no evidence that aIPS stimulation affected the size-weight illusion. Small effects were, however, found on anticipatory force scaling, where grip force was less tuned to object size during initial lifts. These findings suggest that, while aIPS might be peripherally involved in sensorimotor prediction, other brain areas underpin the processes that mediate the size-weight illusion.

## 1 INTRODUCTION

An important prerequisite for skilled object manipulation is the accurate planning of the hand and fingertip movements. This motor plan relies on predictions of object weight from object characteristics, such as its size and material. This anticipatory prediction allows for the generation of fingertip forces scaled to object weight and ensure smooth lifting movements. When the prediction is incorrect, forces can be quickly adjusted (Johansson & Westling, 1988).

The action of lifting an object can also provide more information about the object’s properties, such as its weight. However, weight perception is not always veridical. For instance, perceptual judgements of object heaviness can be influenced by object size, referred to as the size-weight illusion (SWI; Charpentier, 1891). In this illusion, a smaller object is judged to be heavier than an equally-weighted large object (see for reviews Buckingham, 2014; Dijker, 2014; Saccone & Chouinard, 2019). To induce the illusion, differences in the sizes of the objects can be perceived visually, haptically (Buckingham, 2019; Ellis & Lederman, 1993) or even illusory (de Brouwer, Smeets, & Plaisier, 2016) and can be independent of object volume (Plaisier & Smeets, 2015).

Although initially both force scaling and weight perception are influenced by object size, after a few lifts, the force scaling will adapt to the actual weight of the objects, whereas the SWI remains constant (Flanagan & Beltzner, 2000; Flanagan, Bittner, & Johansson, 2008; Grandy & Westwood, 2006). It has been suggested that this is due to sensorimotor memory for object weight, where recent experience with object lifting is used for force scaling of the next lifts (Johansson & Westling, 1988). After considerable practice, the SWI can be diminished as well, but at a much larger timescale (Flanagan et al., 2008). It has therefore been suggested that force scaling and weight perception are underpinned by different sources of information (Flanagan & Beltzner, 2000).

The neural networks for anticipatory force scaling to size and the SWI are largely unknown. Previous research showed that various brain areas are involved in force scaling, when object weight was different than predicted (Jenmalm, Schmitz, Forssberg, & Ehrsson, 2006; Schmitz, Jenmalm, Ehrsson, & Forssberg, 2005). Functional magnetic resonance imaging (fMRI) studies have shown that the posterior parietal cortex is active in force coordination, which plays a crucial role in anticipatory force scaling (Ehrsson, Fagergren, Johansson, & Forssberg, 2003). Using transcranial magnetic stimulation (TMS), the causal role of an area in controlling a specific movement and/or perceptual parameter can be determined. Using this technique, previous research has identified a role of the anterior intraparietal sulcus (aIPS) in force scaling when lifting objects (Dafotakis, Sparing, Eickhoff, Fink, & Nowak, 2008; Davare, Duque, Vandermeeren, Thonnard, & Olivier, 2007). In addition to its role in anticipatory force scaling, it has been shown that aIPS responds to object size in studies with non-human primates (Murata, Gallese, Luppino, Kaseda, & Sakata, 2000) and human brain imaging (Chouinard, Large, Chang, & Goodale, 2009; Monaco, Sedda, Cavina-Pratesi, & Culham, 2015), especially in the context of object grasping (Cavina-Pratesi, Goodale, & Culham, 2007) and relevant grasp dimensions (Monaco et al., 2014). Furthermore, it is known that aIPS is involved in controlling grasping movements in reaching with perturbations to object size (Glover, Miall, & Rushworth, 2005; Tunik, Frey, & Grafton, 2005). Therefore, it seems that aIPS is not only involved in force scaling, but also sensitive to object size. Since object size is often indicative of object weight, this could suggest that aIPS could also be important in anticipatory force scaling to object size.

Regarding the SWI, few neuroimaging studies in healthy subjects have been performed. One functional magnetic resonance imaging (fMRI) study showed that the ventral premotor cortex showed greater levels of adaptation in trials which elicited the SWI compared to when lifting objects of the same size and weight, whereas aIPS mainly responded to size (Chouinard et al., 2009). On the other hand, clinical evidence suggests a role of the posterior parietal cortex in the SWI. A case-study of a patient with lesions in this region had no illusion on the side ipsilateral to the lesion (Li, Randerath, Goldenberg, & Hermsdorfer, 2007), although the same research group found more mixed results in another study (Li, Randerath, Goldenberg, & Hermsdorfer, 2011). Furthermore, it has been demonstrated that object weight could be represented in the lateral occipital cortex (Gallivan, Cant, Goodale, & Flanagan, 2014). This was an interesting result, since this area is also known to be involved in visual as well as haptic object perception (Amedi, Jacobson, Hendler, Malach, & Zohary, 2002; Amedi, Malach, Hendler, Peled, & Zohary, 2001) and especially involved in the representation of object shape (Kourtzi & Kanwisher, 2001). However, it was recently shown that bilateral lesions to the lateral occipital cortex did not affect perception of the SWI in a stroke patient (Buckingham, Holler, Michelakakis, & Snow, 2018), leaving it unclear whether this region has a causal role in weight perception.

To summarize, there is considerable evidence that suggests aIPS could play a role in anticipatory scaling to object size, but its potential contribution to the SWI is less clear. The present study aims to investigate the role of aIPS in force scaling to size and the SWI. Participants lifted objects of different size and weight and judged their heaviness. TMS was used to disrupt aIPS and determine its causal role in motor and perceptual processes. Before performing the task, three groups of participants received theta burst stimulation (Huang, Edwards, Rounis, Bhatia, & Rothwell, 2005) to aIPS, the primary motor cortex (M1) or were delivered a sham condition. Since it is known that force scaling based on sensorimotor memory of object weight is represented in M1 (Chouinard, Leonard, & Paus, 2005), we chose this region as a control area to test whether TMS effects of anticipatory force scaling where not due to general alterations in force control. We expected that stimulation to M1 and aIPS would affect force scaling, where anticipatory force scaling to object size would be influenced by stimulation to aIPS. More specifically, we hypothesized that aIPS stimulation would reduce the difference in planned forces towards small and large objects. In turn, this reduced force scaling to size could also reduce the mismatch between perceived and expected weight, hence leading to a reduced size-weight illusion.

Alternatively, it is also possible to assume that aIPS stimulation, while disrupting object-related motor control processing, would reveal the influence of perceptual processing on the control of force parameters. In other words, the perceptual effects of the SWI may influence force scaling, causing force not to be scaled towards expected, but to perceived object weights. We therefore hypothesized that anticipatory force scaling to object size would be intact (i.e. higher forces for larger object) for initial lifts, but after experiencing the SWI, this scaling would be reduced or even reversed where lower force would be planned for the larger, perceptually lighter, object.

## 2. METHODS

### 2.1 Participants

47 participants (27 female, 22±3.3 years old) took part in the study. They were all right-handed (Edinburgh handedness inventory, mean L.Q: 0.86±0.19 (Oldfield, 1971)) and were screened for potential TMS risks (Rossi, Hallett, Rossini, Pascual-Leone, & Group, 2009). They signed informed consent before participation. Participants were divided into three groups, one for each TMS stimulation site (aIPS: n=16, M1: n=16 or Sham: n=15). Two participants were excluded from force analysis (one from the aIPS group, one from the M1 group), because of technical errors in the force data recording. The study was approved by the medical ethical committee of UZ/KU Leuven.

### 2.2 TMS procedure

Continuous theta burst stimulation (cTBS) was applied before the behavioural task using a 70 mm figure-of-eight TMS DuoMag XT coil (Deymed Diagnostic) or a 70 mm Magstim rapid coil (Magstim). We used a Brainsight system (Rogue Research, Canada) for neuronavigation and recording of electromyography (EMG). EMG was recorded in the right first dorsal interosseus (FDI) using a belly-tendon montage. A ground electrode was placed on the processus styloideus ulnae. To determine the stimulation intensity, we determined the rest (rMT) and active (aMT) motor threshold of FDI when stimulating the motor hotspot, which was defined as the position on M1 that gave the largest motor evoked potential (MEP) in response to TMS. Here, rMT was the stimulation intensity that gave MEPs of at least 50µV in 5/10 stimulations while the hand was at rest and aMT the intensity that gave MEPs in 5/10 trials during contraction of FDI at submaximal levels.

cTBS was applied to the participants following standard procedures (600 pulses, 50 Hz triplets at 5 Hz for a total duration of 40 ms) (Huang et al., 2005). Participants were divided into three groups: aIPS, Sham and M1. The aIPS and M1 group received cTBS at 80% of the aMT over aIPS or M1, respectively. The Sham group received cTBS at a low intensity, namely 40% of aMT over aIPS. Stimulation locations were monitored using Brainsight (Rogue Research) software. In the aIPS group, aIPS was defined anatomically on a structural magnetic resonance imaging (MRI) scan, at the border of the posterior sulcus and the intraparietal sulcus. Brain images were obtained with a 3T scanner (Achieva dstream, Phillips Medical Systems) as high-resolution 3D T1-weighted images (TR=9.7 ms, TE=4.6 ms, field of view = 256×256 mm^2^, 192 slices, voxel size = 0.98 × 0.98 ×1.2 mm^3^). The mean MNI coordinates of aIPS in the aIPS group were (−45±3, −39±6, 48±5). The Sham target and M1 target were defined on a model brain from the Brainsight software, with aIPS MNI coordinates from literature (−43, −39, 46) (Davare, Andres, Clerget, Thonnard, & Olivier, 2007) or the motor hotspot (mean MNI coordinates: −64±14, 2±16, 77±7), respectively. Individual and average stimulation sites are shown in Figure 1C, where all targets are shown projected on the cortex. Note that the coordinates from M1 were recorded from the skin, whereas the aIPS targets were determined on the cortex. In addition, the M1 targets were determined on a model brain, therefore the individual targets are much more variable. In all three groups, the coil was positioned with the handle pointing backwards, roughly 45° from the midline, with a posterior-anterior current direction.

**Figure 1.**
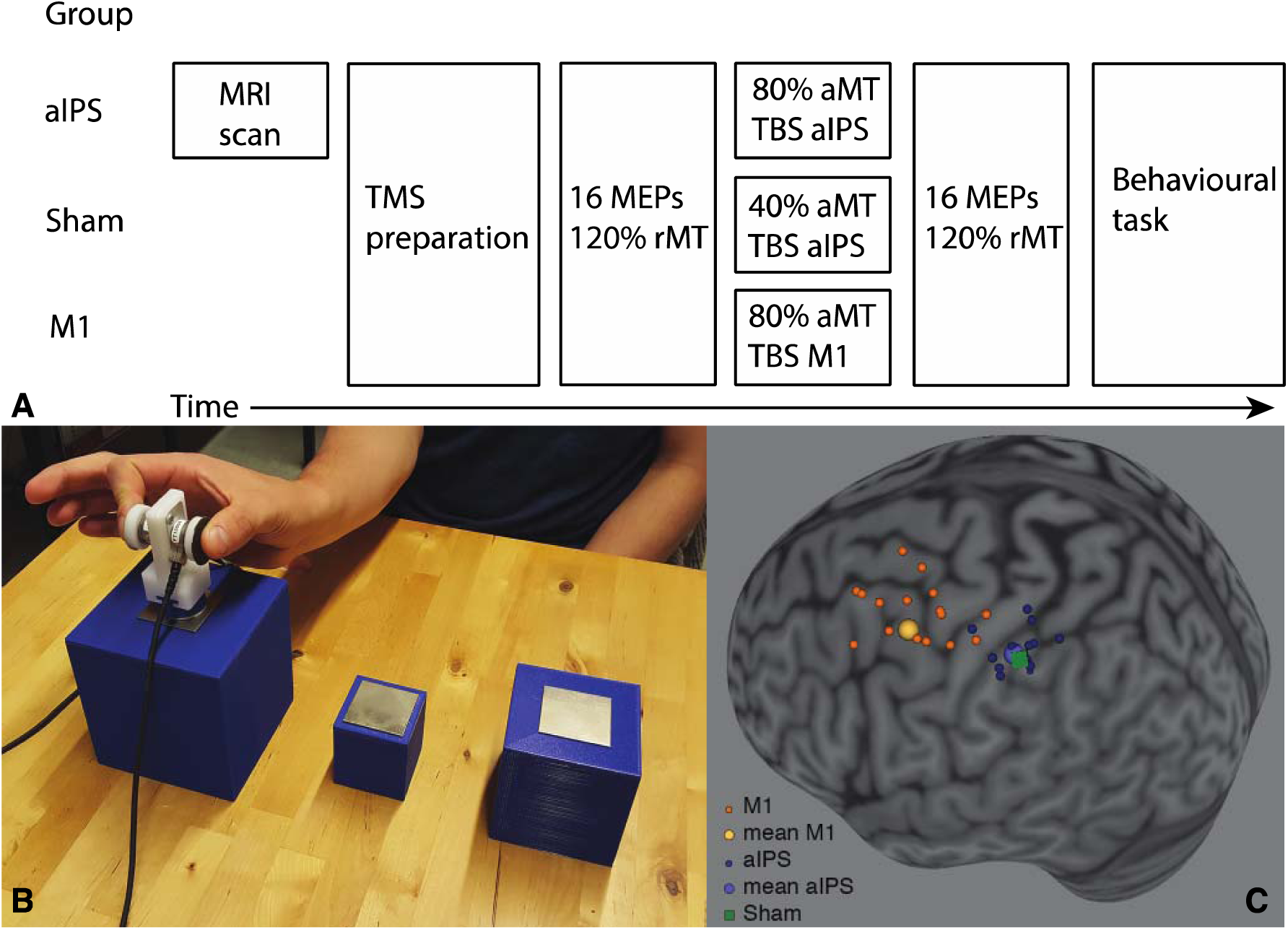
A. Timeline of experiment. Participants were divided into the aIPS, Sham or M1 group. The aIPS group received an MRI scan to anatomically determine the stimulation target. Participants received continuous theta burst stimulation (cTBS) at 40 or 80% of the active motor threshold (aMT). Before cTBS, 16 motor evoked potentials (MEPs) were obtained at 120% rest motor threshold (rMT). After this, participants performed a behavioural task, consisting of lifting objects of different sizes and weights. B. Experimental set-up with large, small and medium sized objects. The medium object was only used in practice trials. For the small and large objects, there was a light and heavy set. A grip manipulandum with force sensors could be quickly attached to the objects using magnets. C. Individual and average stimulation targets, projected on the cortex of the Colin27 MNI reference brain. Note that M1 targets and Sham targets were recorded on a model brain from Brainsight Software.

To measure effects of cTBS on corticospinal excitability, we collected MEPs. Before cTBS, 16 MEPs were collected by stimulating over M1 at an intensity of 120% rMT (pre-MEPs). Next, cTBS was performed after which a 5 min rest period was induced, where the hand did not move. After this rest period, another 16 MEPs were measured with M1 stimulation at 120% rMT (post-MEPs). Following the TMS procedure, participants performed the behavioural task within 45 min after the cTBS. Therefore, the task was completed within the effective period of cTBS as reported in literature (Huang et al., 2005). The timeline of the experiment is illustrated in Figure 1A.

### 2.3 Grasp and Lift Task

Four different 3D-printed cubes were used in the experiment, two large and two small ones. The large cubes measured 10×10×10 cm and the small ones 5×5×5 cm. The objects were filled with lead shot to create two different weights, 190 and 400 g. Therefore, four objects were used: small-light, small-heavy, large-light and large-heavy. A fifth object of a medium size (7.5×7.5×7.5 cm, weight 190 g) was used for practice trials. A pair of force sensors (Nano17, ATI Industrial Automation) was fastened to a 3D-printed manipulandum (Figure 1B). The manipulandum had three magnets in order to quickly attach it to metal squares of 5×5 cm that were glued on top of the objects. The objects were placed behind a screen (Magic Glass) that could switch between a transparent and opaque state.

Participants were instructed to grasp and lift the cube by placing the thumb and index finger on the force sensors. They could lift the cube as soon as the screen turned transparent and the object was visible. Then, they should lift it to a height of approximately 5 cm and hold it in the air until the screen turned opaque again (±3s), upon which they replaced the object on the table. Next, they were asked to give a number best representing the weight of the object on a self-chosen scale. The presentation order of the four objects was pseudo-randomized, in a way that each object size (large or small) followed each object 10 times. Since the first trial was not preceded by any object, one randomly chosen object was added to the trials. This gave a total of 81 trials (4 objects x 2 sizes x 10 orders +1), where each object was presented at least 20 times. To be able to compare the effect of object appearance (i.e. size) before any experience with the objects, the first two lifts were always performed with the small-light and large-light object. Before the start of the experiment, participants performed 10 practice trials with the medium cube.

### 2.4 Data Analysis

#### 2.4.1 MEPs

MEPs were calculated using Brainsight software as the peak-to-peak amplitude of the EMG response. From the 16 MEPs, the first was removed to exclude surprise effects and the remaining 15 were averaged for the pre and post session, respectively. Three MEPs (all in the post-MEP session) were removed because the coil was off-target (>3mm). One participant was excluded from this analysis (M1 group), due to large TMS artefacts in the post-MEP session.

#### 2.4.2 Force and perceptual analysis

Fifteen trials were removed from the perception and force analysis because objects were lifted twice, the object was dropped or touched before the screen turned transparent. Since some of these trials were the first or second trials, 3 participants were excluded from the first-trial analysis (see below) for both the perceptual and force parameters. Two participants were completely excluded (one for aIPS group, one for M1 group) from the force analysis due to technical errors in force data collection. The number of participants used in the data analysis are indicated in all figures.

The weight judgements from participants were converted to z-scores to normalize the values. Forces were filtered with a 2^nd^-order lowpass Butterworth filter with a cut-off frequency of 15 Hz. Grip forces (GF) were defined as the mean of the forces perpendicular to the sensor, load forces (LF) were the sum of the vertical forces. Force rates were the differentiated forces, GFR and LFR, respectively. Parameters of interest were the first peak of GFR (GFR1st), the first peak of LFR (LFR1st) and the loading phase duration (LPD). The first peak of the force rates were the first peaks after grip force onset (GF>0.1 N). To exclude peaks due to noise and positioning the fingers on the sensors, only peaks that were at least 70% of the maximum force rates were included. The load force duration was the time between LF onset (LF>0.1 N) and lift-off (LF>object weight).

#### 2.4.3 Statistics

Paired-samples t-tests were used to determine differences between pre and post-MEPs for each group. Force parameters and perceptual estimates were analysed in three ways. First, we looked at the behavioural effects of size and weight averaged across all lifts for each object. The parameters (GFR1st, LFR1st, LPD, perceptual estimates) obtained for each object were compared in a 2 (object size) x 2 (object weight) x 3 (cTBS group) mixed analysis of variance (ANOVA). Here, object size and object weight were within factors and cTBS group was a between factor.

Secondly, next to effects on the average of all trials, we tested the performance on the first trials only. This was done to test how forces were scaled to object appearance without any previous experience with the object. Therefore, we took the last lift of the practice trials (medium light), the first lift (small light) and the second lift (big light) and compared these in a 3 (size) x 3 (cTBS group) ANOVA. Here, size and cTBS group were within and between factors, respectively. For the perceptual estimates, we compared the first and second lift in a 2 (size) x 3 (cTBS group ANOVA), because no perceptual judgement was obtained for the practice trials with the medium weight. Because for three participants a measurement error was observed in the first or second trial, they were excluded from this analysis.

Finally, we examined the effects of previous lifted weight on current lifted weight. We examined these order effects of object weight since it is known that the weight of previously lifted objects can influence force scaling and weight perception (Johansson & Westling, 1988; van Polanen & Davare, 2015b). Therefore, we ordered the trials in four possible ways: light-light (LL), heavy-light (HL), light-heavy (LH) and heavy-heavy (HH), regardless of object size. For instance, in a heavy-light order, a heavy object was lifted first and the force and perceptual parameters for lifting the light object on the next trial were examined. We conducted a 2 (current weight) x 2 (previous weight) x (cTBS group) ANOVA on the three force parameters and the perceptual estimates. Here, current weight and previous weight were within factors and cTBS group was a between factor.

In case of a main effect or interaction with cTBS group, the ANOVA was split into three separate repeated measures ANOVAs to investigate effects in each cTBS group. If no effect or interaction of cTBS group was found, the groups were pooled into a repeated measures ANOVA to further investigate possible within interaction effects. Further post-hoc tests were performed using t-tests with a Bonferroni correction. A p-value of <0.05 was considered significant.

#### 2.4.4 Bayesian statistics

The absence of a significant effect cannot directly be interpreted as a confirmation of the absence of the effect. It has been suggested that especially in cTBS experiments, additional Bayesian statistics could be helpful (Biel & Friedrich, 2018). Since we were especially interested in effects of object size and the effect of aIPS, we calculated Bayes factor only for the size x cTBS interaction effects. We only did this for the comparison between the aIPS and the Sham condition, where the influence of cTBS over aIPS was compared to the control condition (Sham), specifically for the differences between the two object sizes. We used the methods as described in (Dienes, 2014) and used the free calculator from (Dienes, 2008). In short, the raw interaction effects were calculated as the difference between the object sizes, averaged over object weight, in the Sham condition and the aIPS condition. Next, Bayes factor was determined for these size effects, with as lower limit the effect of the Sham condition and as upper limit the effect of the aIPS condition. These limits follow from the assumption that cTBS over aIPS could maximally completely eradicate the baseline effect in the Sham condition, or, maximally increase the baseline effect with the effect in the aIPS condition. A Bayes factor lower than 1/3 would be interpreted as evidence for the null hypothesis, whereas a Bayes factor higher than 3 would be interpreted as evidence for the alternative (Jeffreys, 1939/1961).

#### 2.4.5 Correlations

To test whether effects on force scaling and perceptual estimates were related, we performed Pearson’s correlations between these parameters. All parameters were converted into z-scores. Then, we performed correlations between effects of size and effects of sensorimotor memory. For the effects of size, we subtracted trials with big objects from small objects for light and heavy objects separately (i.e. small-light – big-light and small-heavy – big-heavy). We correlated these differences for the perceptual estimates with the differences for GFR1st, LFR1st and LPD. The correlations were performed for each cTBS group separately. Similarly, for the sensorimotor memory effects, we subtracted the values for trials that had a previous light object with trials with a previous heavy object for current light and heavy objects separately (i.e. HL-LL and HH-LH). We correlated these differences for the perceptual estimates with those for the three force parameters.

We also performed trial-by-trial correlations for each participant between perceptual estimates and GFR1st and LFR1st. We did this for the light and heavy object set separately, to investigate effects of size not affected by object weight.

## 3 RESULTS

### 3.1 MEPs

Results for the MEPs pre and post cTBS are shown in Figure 2. No significant differences were found between pre and post MEPs for any of the cTBS groups.

**Figure 2.**
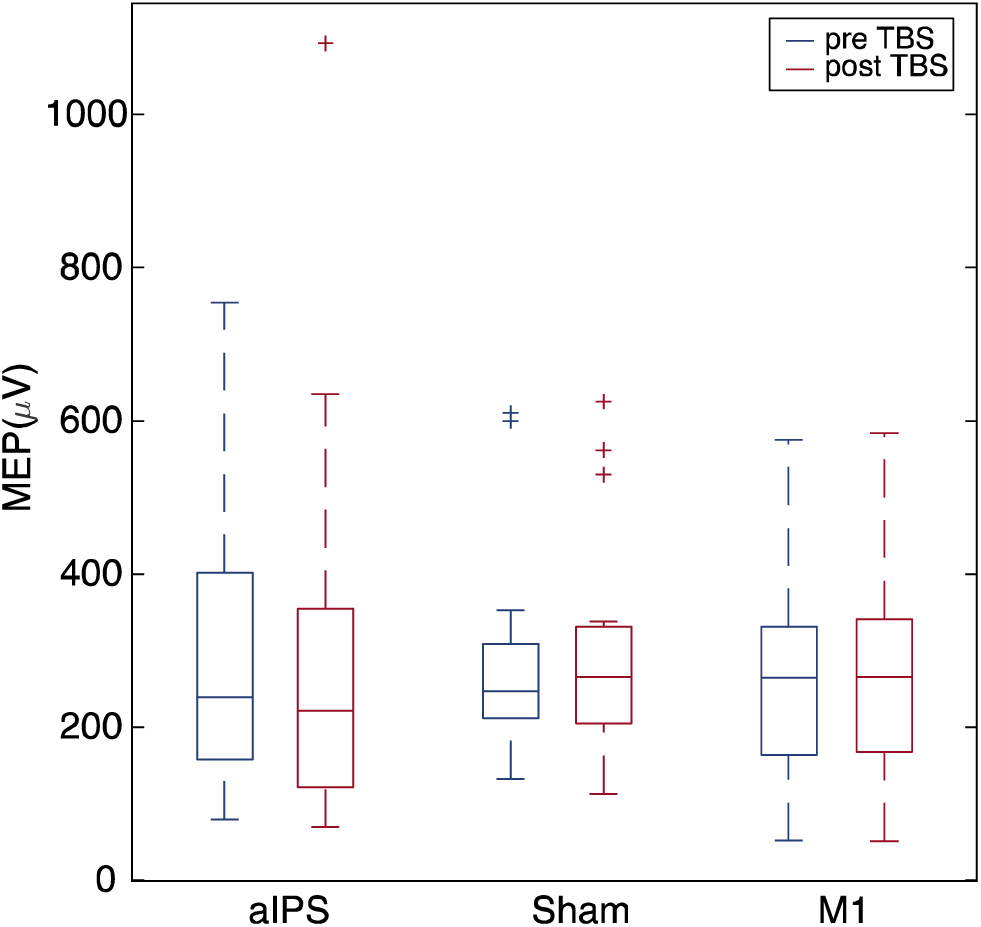
Boxplots for motor evoked potentials (MEP) pre and post continuous theta burst stimulation (cTBS) in each cTBS group (aIPS, N=16; Sham, N=15; M1, N=15). No significant differences were found between pre and post MEPs.

### 3.2 Perceptual estimates were not affected by cTBS

For the normalized perceptual estimates, the judgements were compared for each object (Figure 3). The 2 (mass) × 2 (size) × 3 (cTBS group) ANOVA did not show an effect or interaction with cTBS group. Therefore, a 2 (mass) × 2 (size) ANOVA was performed on the pooled groups. This showed an effect of mass (F(1,46)=5064.3, p<0.001, *η*_p_^2^=0.99), size (F(1,46)=1108.6, p<0.001, *η*_p_^2^=0.96) and an interaction of mass × size (F(1,46)=14.0, p=0.001, *η*_p_^2^=0.23). Post-hoc effects indicated that the light object was perceived to be lighter than the heavy object for both small (p<0.001) and large (p<0.001) sets, as expected in normal weight perception. Furthermore, a size-weight illusion was seen, with small objects perceived as heavier than large objects, both for the light (p<0.001) and heavy (p<0.001) object set. The interaction appears to be explained by a larger-magnitude SWI in heavy objects.

**Figure 3.**
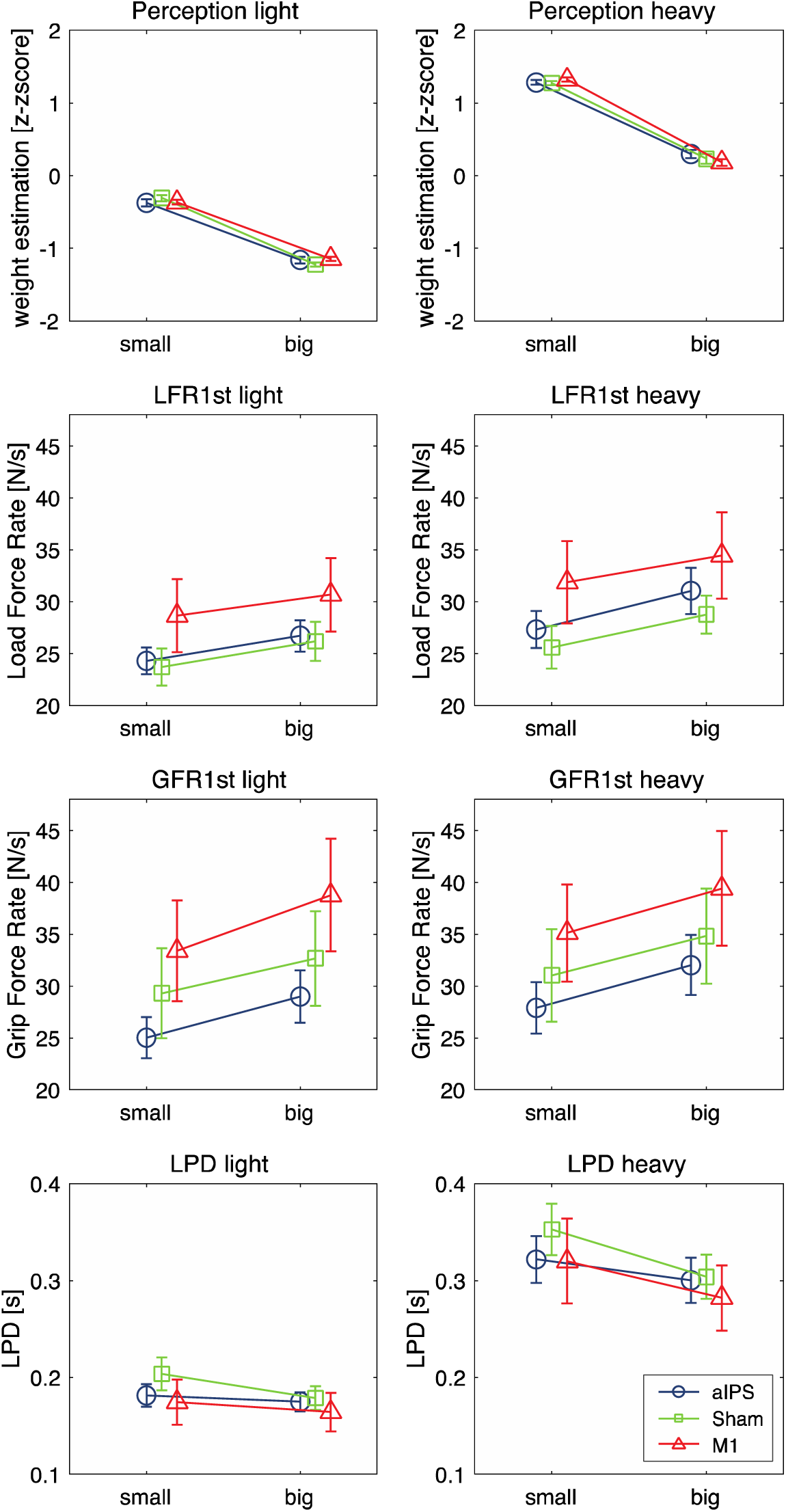
Results for perceptual estimates, first peak of load force rate (LFR1st), first peak of grip force rate (GFR1st) and load force duration (LPD). Values are shown for light (left panels) and heavy objects (right panels), for small and big objects, and for each cTBS group separately (perception: aIPS, N=16; Sham, N=15; M1, N=16. Force parameters: N=15 for all groups). Error bars represent standard errors of the mean. Main effects of size and mass were found for all parameters.

For the first two trials, the two differently-sized objects were also perceived differently (Figure 4). The 2 (size) × 3 (cTBS group) ANOVA revealed an effect of size (F(1,41)=131.7, p<0.001, *η*_p_^2^=0.76). The small object was perceived to be heavier than the large object. There was no effect of cTBS group or an interaction.

**Figure 4.**
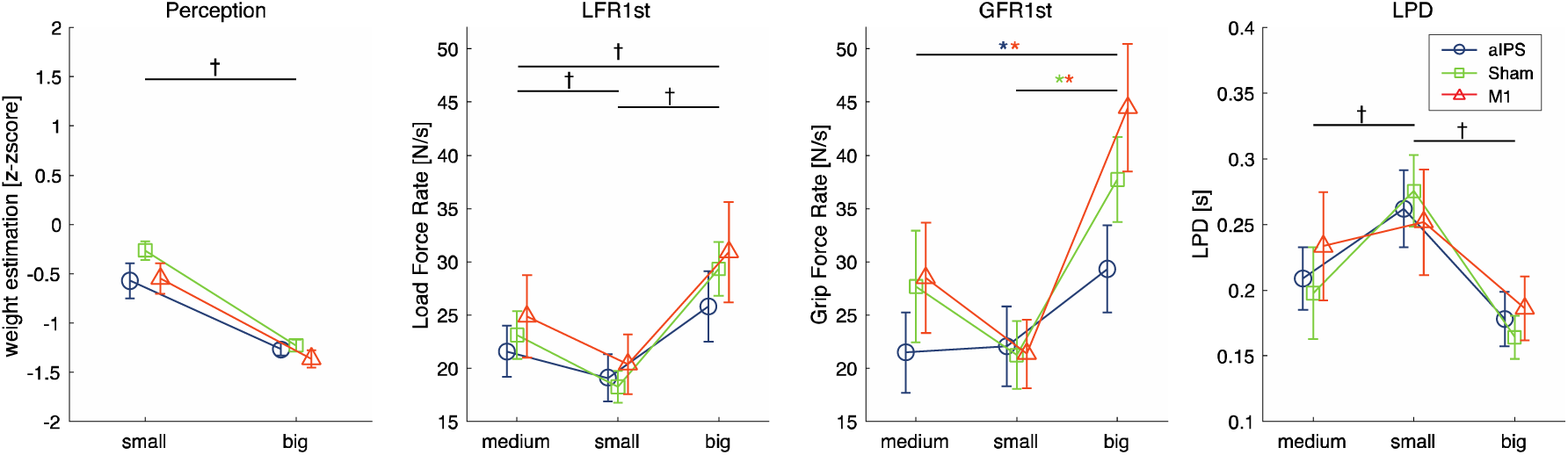
Results for the first two trials (small and big object) for perceptual estimates, first peak of load force rates (LFR1st) and grip force rates (GFR1st) and loading phase durations (LPD). For the force parameters, also the last practice trial (medium object) is shown. Values are shown for each cTBS group (perception: aIPS, N=14; Sham, N=14; M1, N=16. Force parameters: aIPS, N=13; Sham, N=14; M1, N=15). Error bars represent standard error of the mean. †main effect of size. *specific size effect for TBS group.

Finally, we analysed order effects in the perceptual estimates, represented in Figure 5. The 2 (current weight) × 2 (previous weight) × 3 (cTBS group) ANOVA had no effects or interactions with cTBS group. A 2 × 2 ANOVA pooled over the cTBS groups showed main effects of current weight (F(1,46)=5006.0, p<0.001, *η*_p_ ^2^=0.99) and previous weight (F(1,46)=9.3, p=0.004, *η*_p_^2^ =0.17), without an interaction. Again, light objects were perceived as lighter than heavy objects. Furthermore, when a heavy object was previously lifted, objects felt lighter than when a light object was previously lifted.

**Figure 5.**
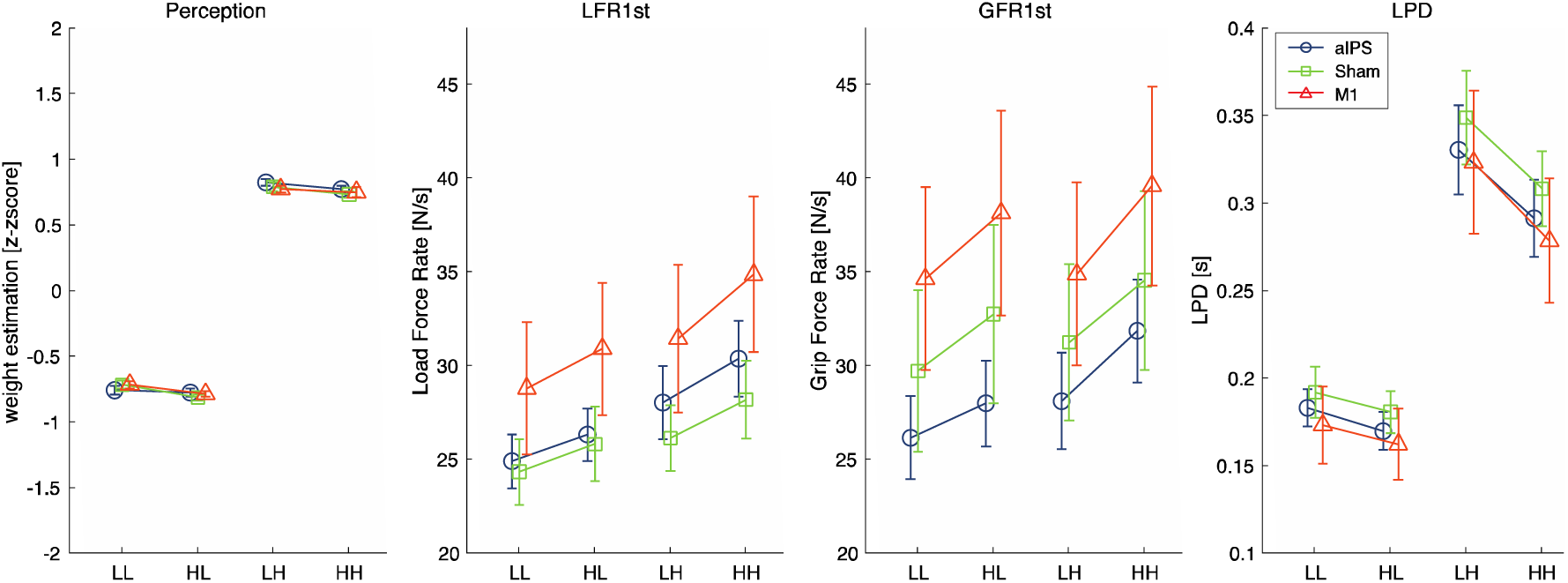
Order effects for perceptual estimates, first peak of load force rates (LFR1st) and grip force rates (GFR1st) and loading phase durations (LPD). Values are shown for the object orders light-light (LL), heavy-light (HL), light-heavy (LH) and heavy-heavy (HH), for each cTBS group (perception: aIPS, N=16; Sham, N=15; M1, N=16. Force parameters: N=15 for all groups). Error bars represent standard error of the mean. Effects of previous and current weight were found for all parameters.

These findings indicate that participants experienced a size-weight illusion, already from the first trial, and replicate the perceptual bias found in (van Polanen & Davare, 2015b). However, these perceptual estimates were not affected by cTBS.

### 3.3 Force parameters

#### 3.3.1 Objects were not lifted differently in response to cTBS

Results for the force parameters are shown in Figure 3. In the analysis for all objects, no main effects of cTBS group were found for LFR1st, GFR1st or LPD, so 2 (mass) × 2 (size) ANOVAs were performed on the pooled data. For LFR1st, main effects of mass (F(1,44)=58.2, p<0.001, *η*_p_^2^=0.57) and size (F(1,44)=54.2, p<0.001, *η*_p_^2^=0.55) were found, where force rates were higher for heavy and large objects compared to light and small objects, respectively.

For GFR1st, the ANOVA revealed an effect of mass (F(1,44)=43.3, p<0.001, *η* _p_ ^2^=0.50), where GFR1st was higher for heavy objects compared to light objects. Furthermore, an effect of size (F(1,44)=51.3, p<0.001, p2=0.54) showed that GFR1st was higher for larger than smaller objects. It must be noted that for GFR1st, a trend toward an interaction between mass and cTBS group was observed (F(2,44)=3.0, p=0.061, *η*p^2^=0.13). However, separate 2 (mass) × 2 (size) ANOVA’s revealed main effects of mass and size for each cTBS group and no significant differences between the cTBS groups were observed for either light or heavy objects. Therefore, no obvious differences between the cTBS groups were present.

For LPD, main effects of mass (F(1,44)=223.6, p<0.001, *η*_p_^2^=0.84), size (F(1,44)=26.8, p<0.001, *η*_p_^2^=0.38) and an interaction of mass × size (F(1,44)=14.4, p<0.001, *η*_p_ ^2^=0.25) were found. Post-hoc tests indicated that the light objects were lifted with shorter LPDs than heavy objects, both for small (p<0.001) and large (p<0.001) objects. In addition, the smaller objects were lifted with longer LPDs than the large objects, both in the light (p=0.020) and heavy (p<0.001) object set. The interaction did not reveal new insights, but might be explained by larger effects of size for heavy objects.

In sum, it seemed that force scaling was adapted to object size and mass, but this was not affected by cTBS.

#### 3.3.2 cTBS influenced grip force scaling when lifting without object experience

The results for the last practice trial and the first two trials are shown in Figure 4. For LFR1st, the first trials were only affected by size (F(2,78)=26.4, p<0.001, *η*_p_ ^2^=0.40), with no main effect or interaction with cTBS group. Post-hoc analyses showed that all sizes differed, where large objects were being lifted with higher force rates compared to small (p<0.001) and medium sizes (p=0.001), and the medium object was lifted with a higher LFR1st than the small one (p=0.004).

For GFR1st in the first trials, the 3 (size) × 3 (cTBS group) demonstrated a main effect of size (F(2,78)=39.4, p<0.001, *η*_p_^2^ =0.50), but also an interaction of size × cTBS group (F(4,78)=3.3, p=0.014, *η*_p_ ^2^=0.15). Therefore, we performed separate ANOVAs for each cTBS group. We found effects of size in each cTBS group (aIPS: F(2,24)=6.3, p=0.007, _p_ ^2^=0.34; M1: F(2,28)=28.4, p<0.001, *η*_p_^2^ =0.67; Sham: F(2,26)=10.6, p<0.001, *η*_p_^2^ =0.45). However, post-hoc analysis showed different results. In the aIPS group, only a difference between the medium and large object was found (p=0.027), where GFR1st was larger for the large object. No significant differences were found between small and medium (p=1.00) or small and big (p=0.053) objects. In the M1 group, the large object had a higher GFR1st compared to both the small (p<0.001) and medium (p<0.001) object. Small and medium objects were not significantly different (p=0.062). Finally, with Sham stimulation, the GFR1st differed only between the small and large object (p=0.001), with a larger GFR1st for the large object. Differences between small and medium (p=0.161) or medium and large objects (p=0.12) were not significant. No differences between the cTBS groups were found for any of the object sizes.

For LPD, only a main effect of size (F(2,78)=15.2, p<0.001, *η*_p_^2^ =0.28) was found, where small objects had shorter LPDs than medium (p=0.012) and large (p<0.001) objects.

To summarize, on the first trials forces were already scaled towards object size. When cTBS was applied, this affected the grip force scaling towards object size differently in the aIPS, M1 and Sham group. Specifically, no difference in grip force rate between small and large objects was found after cTBS on aIPS.

#### 3.3.3 Force scaling according to previous object weight was largely unaffected by cTBS

Effects of object weight order are shown in Figure 5. The analysis on object weight order on the GFR1st revealed main effects of current weight (F(1,42)=36.5, p<0.001, *η*_p_^2^ =0.47), previous weight (F(1,42)=76.6, p<0.001, *η*_p_^2^ *η*_p_^2^ =0.65), but also interactions of previous weight × current weight (F(1,42)=4.1, p=0.049, *η*_p_ ^2^=0.09) and current weight × cTBS group (F(2,42)=4.0, p=0.025, *η*_p_ ^2^=0.16). To analyse the interaction with cTBS group, we performed separate 2 × 2 ANOVAs for each cTBS group. For all cTBS groups, a main effect of current weight was found (aIPS: F(1,14)=21.4, p<0.001, *η*_p_ ^2^=0.61; M1: F(1,14)=5.1, p=0.041, *η*_p_ ^2^=0.27; Sham: F(1,14)=10.3, p=0.006, *η*_p_^2^=0.42), where the light objects had a lower GFR1st than heavy objects. In addition, for all cTBS groups, a main effect of previous weight was found (aIPS: F(1,14)=34.9, p<0.001, *η*_p_ ^2^=0.71; M1: F(1,14)=34.3, p<0.001, *η*_p_ ^2^=0.71; Sham: F(1,14)=16.5, p=0.001, *η*_p_ ^2^=0.54). Here, a previous light weight resulted in a lower GFR1st than a previous heavy weight. No interaction between current and previous weight was found in any cTBS group. Also, no differences between the cTBS groups were found for light nor heavy current weights. Therefore, similar to the trend of the interaction of weight and cTBS group for the object analysis (see above), this interaction between current weight and cTBS group is difficult to explain. From Figure 5, it appears that the difference between light and heavy objects is slightly reduced after cTBS on M1 and increased after cTBS on aIPS.

For the LFR1st and LPD, no main or interaction effects with cTBS group were found. Therefore, the data was pooled over cTBS group in a 2 × 2 ANOVA. For LFR1st, main effects of current weight (F(1,44)=53.2, p<0.001, *η*_p_^2^=0.55), previous weight (F(1,44)=46.6, p<0.001, *η*_p_^2^=0.51) and an interaction of current weight × previous weight were found (F(1,44)=4.9, p=0.031, *η*_p_ ^2^=0.10). Post-hoc analysis indicated that LFR1st was higher for current heavy than current light objects, both when previously a heavy (p<0.001) or light objects (p<0.001) was lifted. Furthermore, if previously a heavy object was lifted, LFR1st was higher compared to previously lifting a light object, both for current light (p<0.001) and heavy objects (p<0.001). For LPD, main effects of current weight (F(1,44)=234.2, p<0.001, *η*_p_^2^=0.84), previous weight (F(1,44)=80.7, p<0.001, *η*_p_^2^=0.65) and an interaction of current weight × previous weight (F(1,44)=28.1, p<0.001, *η*_p_^2^=0.39) were found. Post-hoc analyses revealed that the LPD was shorter for lifting light objects compared to heavy ones, both when the previous object was light (p<0.001) or heavy (p<0.001). When the previous object was light, LPDs were longer, both when the current object was light (p<0.001) and heavy (p<0.001). The interactions did not reveal new insights, but could result from slightly larger order effects for heavy objects than light ones.

In sum, force scaling did also depend on the weight of the previous lifted object. However, this effect was not influenced by cTBS.

### 3.4 Correlations between force parameters and perceptual estimates

To test whether effects of object size on the force scaling and perceptual ratings were related, we calculated correlations of the size effects on force parameters and perceptual estimates. Here, we found that in the Sham condition, there were significant correlations, but only for heavy objects. These results are shown in Figure 6. A negative relation was found between LFR1st and perception (R=-0.58, p=0.023) and between LPD and perception (R=0.56, p=0.029). The correlation of GFR1st and perception was almost significant (R=-0.51, p=0.051). These relations indicate that the amount of scaling to size in force parameters is related to the strength of the SWI. By contrast, no significant correlations were found for the cTBS conditions.

**Figure 6.**
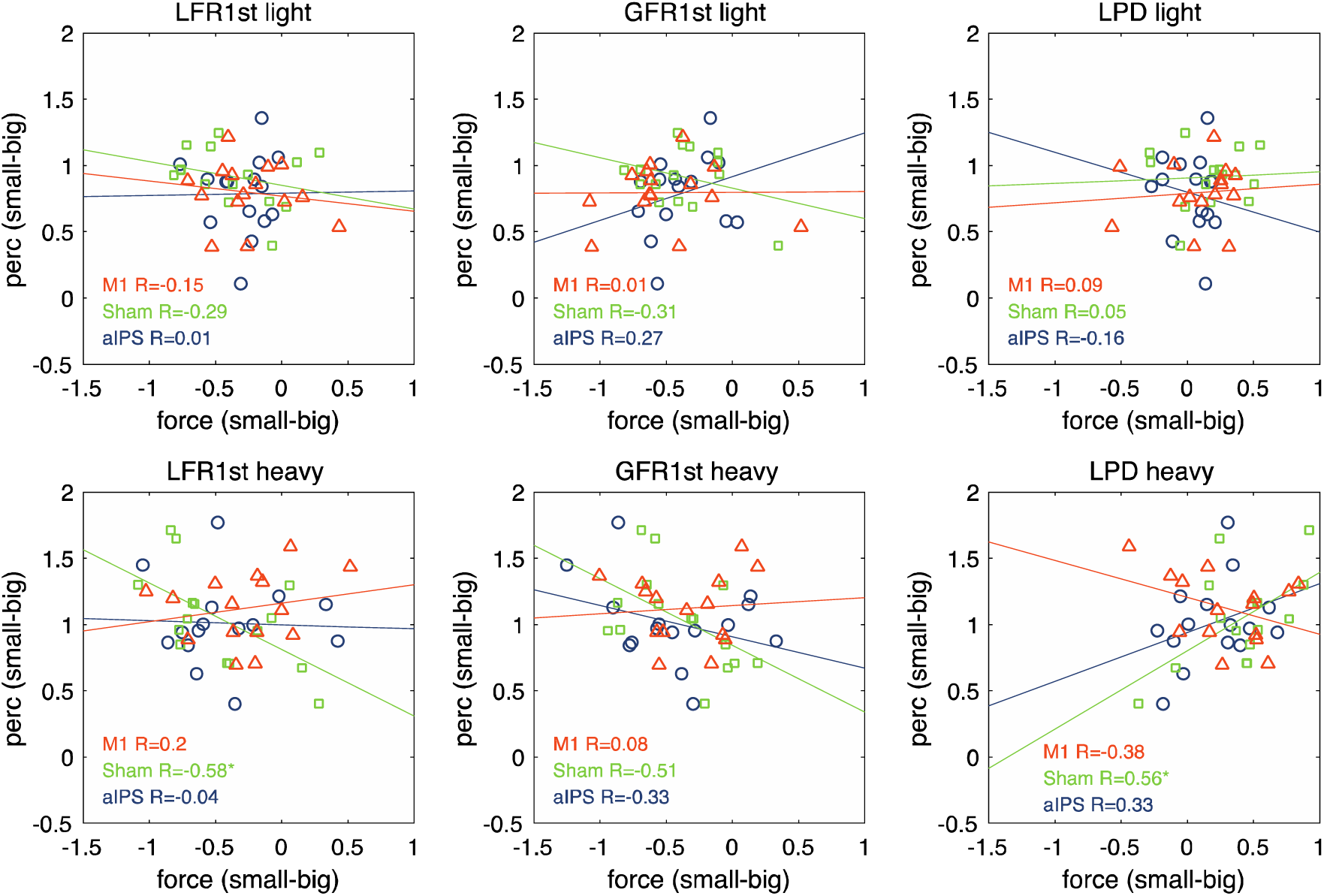
Correlations between size effects of perception and size effects on force parameters first peak of load force rate (LFR1st) and grip force rate (GFR1st) and loading phase duration (LPD). Values are shown for light (top panels) and heavy (bottom panels), for each cTBS group separately (all N=15). *p<0.05

Average trial by trial correlations are shown in Table 1, where most R values were significantly different from zero. Since the correlations were performed separately between object weights, variations in values will most likely reflect variations due to object size. Hence, a significant correlation suggests a relation between force scaling to size and SWI strength. However, no apparent differences were visible between the cTBS groups.

**Table 1.**
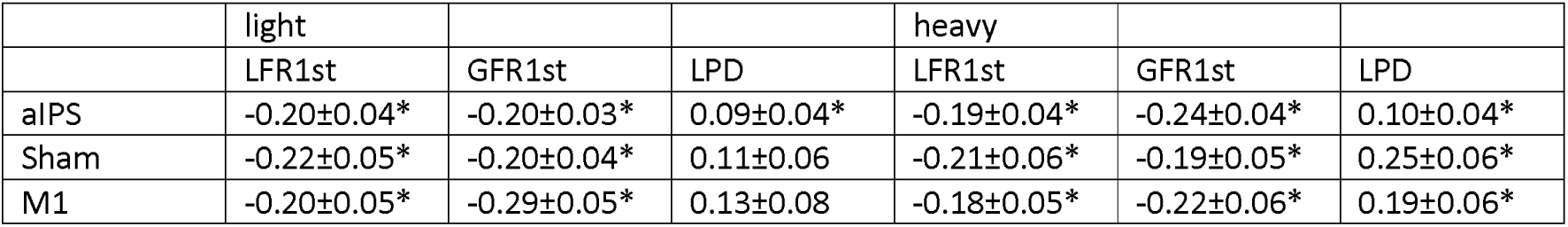
Mean R values for trial by trial correlations between perceptual estimates and force scaling parameters (first peak of load force rate, LFR1st; first peak of grip force rate, GFR1st; and load force duration, LPD) for light and heavy object separately. Values indicate mean±standard error. *p<0.05 significant from zero with one-sample t-test.

|  | light |  |  | heavy |  |  |
| --- | --- | --- | --- | --- | --- | --- |
|  | LFR1st | GFR1st | LPD | LFR1st | GFR1st | LPD |
| aIPS | -0.20±0.04* | -0.20±0.03* | 0.09±0.04* | -0.19±0.04* | -0.24±0.04* | 0.10±0.04* |
| Sham | -0.22±0.05* | -0.20±0.04* | 0.11±0.06 | -0.21±0.06* | -0.19±0.05* | 0.25±0.06* |
| M1 | -0.20±0.05* | -0.29±0.05* | 0.13±0.08 | -0.18±0.05* | -0.22±0.06* | 0.19±0.06* |

Furthermore, to test whether the order of object weight similarly affected force parameters and perceptual estimations, we examined whether the sensorimotor memory effects correlated with order effects on the perceptual estimates; none of these correlations were significant.

### 3.5 Bayesian statistics on size x cTBS interaction

To determine the Bayes Factor for the size × cTBS interaction, we first determined the F-value of this effect for aIPS and Sham only. We performed a 2 (mass) × 2 (size) × 2 (cTBS) mixed ANOVA on the perceptual z-scores, GFR1st, LFR1st and LPD. We found the following results for the size x cTBS interaction: perception (F(1,26)=1.5, p=0.229, *η*_p_^2^=0.05), GFR1st (F(1, 26)=0.2, p=0.691, *η*_p_ ^2^=0.01), LFR1st (F(1, 26)=0.1, p=0.741, *η*_p_ ^2^=0.00) and LPD (F(1, 26)=5.0, p=0.034, *η*_p_ ^2^=0.15). Note that the effect was significant for LPD, but not for the other variables. This resulted in Bayes Factors of 0.21 for perception, 0.32 for LFR1st, 0.42 for GFR1st and 5.55 for LPD. These values indicate evidence for the null-hypothesis for perceptual estimates. Note that although GFR1st and LFR1st Bayes Factor suggests inconclusive evidence, the value is very close to 1/3. By contrast, for LPD, there is substantial evidence for the alternative hypothesis, indicating an effect of cTBS over aIPS on anticipatory force scaling to size.

Furthermore, we also performed this analysis on the data of the first two trials. This represents the effect of object size (small or heavy) and cTBS (aIPS vs Sham). A new 2 (size) x 2 (cTBS) mixed ANOVA on the first two trials gave the following results: perception (F(1,0)=2.1, p=0.155, *η*_p_^2^ =0.08), GFR1st (F(1,0)=4.8, p=0.038, *η*_p_^2^ =0.16), LFR1st (F(1,0)=2.6, p=0.119, *η*_p_^2^ =0.09) and LPD (F(1,0)=1.2, p=0.280, *η*_p_^2^=0.04). Note that the effect was only significant in GFR1st, also found in the original analysis. The resulting Bayes Factors were 0.80 for perception, 1.40 for LFR1st, 4.71 for GFR1st and 0.59 for LPD. For GFR1st this gave substantial evidence for the alternative hypothesis, confirming the significant p-value. For the other variables, inconclusive evidence was found.

## 4 DISCUSSION

The aim of the present study was to evaluate the role of the anterior intraparietal sulcus (aIPS) in anticipatory force scaling to object size and the size-weight illusion (SWI). We used non-invasive brain stimulation to disrupt aIPS prior to a behavioural task. More specifically, continuous theta burst stimulation (cTBS) was applied over aIPS and compared to two control conditions, where cTBS was either applied over the primary motor cortex (M1) or a sham stimulation was performed. In the behavioural task, participants lifted objects of different sizes and weights while their fingertip forces were measured and after each lift they reported felt object heaviness. We found no effect of aIPS stimulation on the SWI and only minor effects on force scaling. This suggest that aIPS is unlikely to be causally involved in the SWI, and does not appear to play a primary role in anticipatory force scaling based on visual cues to object size.

A robust SWI was observed in our experiments, with smaller objects perceived to be heavier than large objects of the same mass, in both the light and heavy object set. The SWI effect was not affected by cTBS applied over aIPS nor M1. To further affirm these non-significant results, we calculated Bayes Factors and these indicated support for the null hypothesis. Therefore, we conclude that aIPS does not play a role in the SWI. Previous clinical evidence suggested a possible role for the posterior parietal cortex, but results were mixed (Li et al., 2007, 2011). It seems likely instead that other areas are more involved in the SWI, such as the ventral premotor cortex (Chouinard et al., 2009) or the lateral occipital cortex (Gallivan et al., 2014), although lesions this latter area did not seem to affect the SWI (Buckingham et al., 2018).

When lifting objects of different sizes, participants scaled their forces to object size, with higher rates of force used to lift large objects compared to small objects. We found this for the first trials, but surprisingly this behaviour was maintained over the course of the experiment. Previous research showed that forces adapted to actual object weight after some experience with the objects (Buckingham & Goodale, 2010; Flanagan & Beltzner, 2000; Flanagan et al., 2008). However, in these earlier studies objects all had the same weight and were presented in alternating order. An earlier study indicated that a random presentation can induce force scaling to object size compared to lifting object in a consecutive order (Gordon, Forssberg, Johansson, & Westling, 1991). In the present experiment, two sets of object weight were presented in a random order. Participants could not use earlier experience with the objects to accurately predict object weight, since an object of a specific size could be either light or heavy and, therefore, they might still have relied on size priors to scale their forces.

Contrary to our hypothesis, we did not find large effects of stimulation over aIPS on anticipatory force scaling to size. Grip force scaling was slightly reduced in the first trials and the loading phase duration appeared to be slightly altered, but overall effects were minor. Although aIPS plays a role in force scaling (Dafotakis et al., 2008; Davare, Andres, et al., 2007), it might not be involved in anticipatory force scaling to object size. It is likely that aIPS is more concerned with online control and error corrections, such as known for grasp parameters (Cavina-Pratesi et al., 2007; Cohen, Cross, Tunik, Grafton, & Culham, 2009; Davare, Rothwell, & Lemon, 2010; Rice, Tunik, & Grafton, 2006) and less with anticipatory force scaling to object properties. Considering the large network of areas involved in planning and control of grasping behaviour (Grafton, 2010), anticipatory scaling could be governed by other areas, such as the premotor cortex (Chouinard et al., 2005; Dafotakis et al., 2008; van Nuenen, Kuhtz-Buschbeck, Schulz, Bloem, & Siebner, 2012).

Several previous studies showed that fingertip forces and perceptual estimates adapt differently to repeated object lifting with SWI objects (Chang, Flanagan, & Goodale, 2008; Flanagan & Beltzner, 2000; Flanagan et al., 2008; Grandy & Westwood, 2006; Trewartha & Flanagan, 2017), where the illusion is still present after several lifts but the forces are scaled correctly for the equally weighting objects. Interestingly, we did not only find that the anticipatory scaling to object size remained when objects were presented randomly, but we also found correlations with perceptual effects. That is, the effect of size on force scaling was related to the magnitude of the SWI. However, these effects should be interpreted with caution, since we only observed small correlations in trial by trial comparisons and only in heavy objects for between-subject correlations. Furthermore, the between-subject correlations disappeared both after stimulation on M1 and aIPS, but there appeared to be no difference between cTBS conditions when correlating individual trials. The disappearance of the relation between force and perceptual measures when stimulating motor-related areas is of interest, but our dataset is limited to draw strong conclusions from these results. Although this relation requires further research, it is noteworthy that Gordon et al. (1991) observed that participants who did not show an SWI, also showed a probing strategy with little force scaling to size, further suggesting links between lifting behaviour and perceptual estimates.

This potential relationship between perceptual and action processes could be linked to interactions between the dorsal and ventral stream which must underpin object interaction behaviour (Goodale & Milner, 1992; Milner & Goodale, 2008). In this theory, the dorsal stream is involved with processing action-related information, whereas the ventral stream is concerned with perceptual processes, such as object recognition. However, it has been acknowledged that these streams also interact (see for reviews e.g. Cloutman, 2013; van Polanen & Davare, 2015a). Since aIPS is part of the dorsal stream, it is possible that disrupting this area could allow for more ventral stream influence on action processes. We hypothesized that the perceptual effect of the SWI would influence force scaling, where the perceptual light large object would be lifted with a lower force instead of the initial high force. However, it seemed that effects of aIPS stimulation were most prominent in the first trials. Although we found small decreases of grip forces for the large object, this was actually most apparent on the first trial, when participants did not have any perceptual experience with the objects. Therefore, it seems unlikely that alterations in force scaling would be caused by influence of ventral stream processes and perceptual object experience of the SWI.

Besides the limited effects of aIPS stimulation on force scaling, we also did not find effects of M1 stimulation. The MEPs were not altered after cTBS. This is in contrast to previous literature that found that MEPs decreased after cTBS over M1 (Huang et al., 2005). However, it has been acknowledged that the effects can be very variable (Hamada, Murase, Hasan, Balaratnam, & Rothwell, 2013; Lopez-Alonso, Cheeran, Rio-Rodriguez, & Fernandez-Del-Olmo, 2014). It must also be noted that the MEPs in this study were very small, leaving little room for decrements. For example, in de study of Huang et al. (2005), the baseline was set at the 1mV threshold, whereas we used 120% of rMT. In our case, this led to baseline values below 1mV, which might have not been decreased visibly by cTBS. Furthermore, stimulation of M1 did not affect sensorimotor memory, whereas this was shown in previous studies (Chouinard et al., 2005; Nowak, Voss, Huang, Wolpert, & Rothwell, 2005). However, since objects also varied in size, it is possible that sensorimotor memory effects were weaker, as forces were also scaled to object size and not only to previous experienced weight. With more variability in force parameters, small effects of M1 stimulation on sensorimotor memory might have been masked.

To conclude, we investigated the role of aIPS in force scaling to size and the SWI. While aIPS might play a minor role in force scaling to size, it does not seem to be involved in mediating the SWI.

## ACKNOWLEDGEMENTS

This research was supported by Fonds Wetenschappelijk Onderzoek grants to VVP (FWO post-doctoral fellowship, Belgium, 12×7118N) and MD (FWO Odysseus, Belgium, G/0C51/13N). The authors would like to thank members and students from the motor control and neuroplasticity lab for their help in data collection: L. Pauwels, L. Hermans, K. Heise, K. Vandevoorde, M. Gann, G. Rens, P. Verstraete and I. Meeusen.

## AUTHOR CONTRIBUTIONS

Conceptualization: VvP, GB, MD. Investigation: VvP. Formal analysis: VvP. Writing – Original draft: VvP. Visualization: VvP. Writing – Review & Editing: VvP, GB, MD.

## REFERENCES

Amedi, A., Jacobson, G., Hendler, T., Malach, R., & Zohary, E. (2002). Convergence of visual and tactile shape processing in the human lateral occipital complex. Cerebral Cortex, 12(11), 1202–1212. doi: 10.1093/cercor/12.11.1202

Amedi, A., Malach, R., Hendler, T., Peled, S., & Zohary, E. (2001). Visuo-haptic object-related activation in the ventral visual pathway. Nature Neuroscience, 4(3), 324–330. doi: 10.1038/85201

Biel, A. L., & Friedrich, E. V. C. (2018). Why You Should Report Bayes Factors in Your Transcranial Brain Stimulation Studies. Front Psychol, 9(1125), 1125. doi: 10.3389/fpsyg.2018.01125

Buckingham, G. (2014). Getting a grip on heaviness perception: a review of weight illusions and their probable causes. Experimental Brain Research, 232(6), 1623–1629. doi: 10.1007/s00221-014-3926-9

Buckingham, G. (2019). Examining the size-weight illusion with visuo-haptic conflict in immersive virtual reality. Q J Exp Psychol (Hove), 72(9), 2168–2175. doi: 10.1177/1747021819835808

Buckingham, G., & Goodale, M. A. (2010). The influence of competing perceptual and motor priors in the context of the size-weight illusion. Experimental Brain Research, 205(2), 283–288. doi: 10.1007/s00221-010-2353-9

Buckingham, G., Holler, D., Michelakakis, E. E., & Snow, J. C. (2018). Preserved Object Weight Processing after Bilateral Lateral Occipital Complex Lesions. J Cogn Neurosci, 30(11), 1683–1690. doi: 10.1162/jocn_a_01314

Cavina-Pratesi, C., Goodale, M. a., & Culham, J. C. (2007). FMRI reveals a dissociation between grasping and perceiving the size of real 3D objects. PLoS ONE, 2(5), e424. doi: 10.1371/journal.pone.0000424

Chang, E. C., Flanagan, J. R., & Goodale, M. A. (2008). The intermanual transfer of anticipatory force control in precision grip lifting is not influenced by the perception of weight. Experimental Brain Research, 185, 319–329. doi: 10.1007/s00221-007-1156-0

Charpentier, A. (1891). Analyse expérimentale quelques éléments de la sensation de poids. Archives de Physiologie Normales et Pathologiques, 3, 122–135.

Chouinard, P. A., Large, M. E., Chang, E. C., & Goodale, M. A. (2009). Dissociable neural mechanisms for determining the perceived heaviness of objects and the predicted weight of objects during lifting: an fMRI investigation of the size-weight illusion. NeuroImage, 44(1), 200–212. doi: 10.1016/j.neuroimage.2008.08.023

Chouinard, P. A., Leonard, G., & Paus, T. (2005). Role of the primary motor and dorsal premotor cortices in the anticipation of forces during object lifting. Journal of Neuroscience, 25(9), 2277–2284. doi: 10.1523/JNEUROSCI.4649-04.2005

Cloutman, L. L. (2013). Interaction between dorsal and ventral processing streams: where, when and how? Brain Lang, 127(2), 251–263. doi: 10.1016/j.bandl.2012.08.003

Cohen, N. R., Cross, E. S., Tunik, E., Grafton, S. T., & Culham, J. C. (2009). Ventral and dorsal stream contributions to the online control of immediate and delayed grasping: a TMS approach. Neuropsychologia, 47(6), 1553–1562. doi: 10.1016/j.neuropsychologia.2008.12.034

Dafotakis, M., Sparing, R., Eickhoff, S. B., Fink, G. R., & Nowak, D. A. (2008). On the role of the ventral premotor cortex and anterior intraparietal area for predictive and reactive scaling of grip force. Brain Research, 1228, 73–80. doi: 10.1016/j.brainres.2008.06.027

Davare, M., Andres, M., Clerget, E., Thonnard, J. L., & Olivier, E. (2007). Temporal dissociation between hand shaping and grip force scaling in the anterior intraparietal area. Journal of Neuroscience, 27(15), 3974–3980. doi: 10.1523/JNEUROSCI.0426-07.2007

Davare, M., Duque, J., Vandermeeren, Y., Thonnard, J. L., & Olivier, E. (2007). Role of the ipsilateral primary motor cortex in controlling the timing of hand muscle recruitment. Cerebral Cortex, 17(2), 353–362. doi: 10.1093/cercor/bhj152

Davare, M., Rothwell, J. C., & Lemon, R. N. (2010). Causal connectivity between the human anterior intraparietal area and premotor cortex during grasp. Current Biology, 20(2), 176–181. doi: 10.1016/j.cub.2009.11.063

de Brouwer, A. J., Smeets, J. B., & Plaisier, M. A. (2016). How Heavy Is an Illusory Length? Iperception, 7(5), 2041669516669155. doi: 10.1177/2041669516669155

Dienes, Z. (2008). Understanding psychology as a science: An introduction to scientific and statistical inference: Macmillan International Higher Education.

Dienes, Z. (2014). Using Bayes to get the most out of non-significant results. Front Psychol, 5(781), 781. doi: 10.3389/fpsyg.2014.00781

Dijker, A. J. (2014). The role of expectancies in the size-weight illusion: a review of theoretical and empirical arguments and a new explanation. Psychon Bull Rev, 21(6), 1404–1414. doi: 10.3758/s13423-014-0634-1

Ehrsson, H. H., Fagergren, A., Johansson, R. S., & Forssberg, H. (2003). Evidence for the involvement of the posterior parietal cortex in coordination of fingertip forces for grasp stability in manipulation. Journal of Neurophysiology, 90(5), 2978–2986. doi: 10.1152/jn.00958.2002

Ellis, R. R., & Lederman, S. J. (1993). The role of haptic versus visual volume cues in the size-weight illusion. Percept Psychophys, 53(3), 315–324. doi: 10.3758/bf03205186

Flanagan, J. R., & Beltzner, M. A. (2000). Independence of perceptual and sensorimotor predictions in the size-weight illusion. Nature Neuroscience, 3(7), 737–741. doi: 10.1038/76701

Flanagan, J. R., Bittner, J. P., & Johansson, R. S. (2008). Experience can change distinct size-weight priors engaged in lifting objects and judging their weights. Current Biology, 18(22), 1742–1747. doi: 10.1016/j.cub.2008.09.042

Gallivan, J. P., Cant, J. S., Goodale, M. A., & Flanagan, J. R. (2014). Representation of object weight in human ventral visual cortex. Current Biology, 24(16), 1866–1873. doi: 10.1016/j.cub.2014.06.046

Glover, S., Miall, R. C., & Rushworth, M. F. (2005). Parietal rTMS disrupts the initiation but not the execution of on-line adjustments to a perturbation of object size. Journal of Cognitive Neuroscience, 17(1), 124–136.

Goodale, M. A., & Milner, A. D. (1992). Seperate visual pathways for perception and action. Trends in Neurosciences, 15, 20–25.

Gordon, A. M., Forssberg, H., Johansson, R. S., & Westling, G. (1991). The integration of haptically acquired size information in the programming of precision grip. Experimental Brain Research, 83(3), 483–488.

Grafton, S. T. (2010). The cognitive neuroscience of prehension: recent developments. Experimental Brain Research, 204(4), 475–491. doi: 10.1007/s00221-010-2315-2

Grandy, M. S., & Westwood, D. A. (2006). Opposite perceptual and sensorimotor responses to a size-weight illusion. Journal of Neurophysiology, 95(6), 3887–3892. doi: 10.1152/jn.00851.2005

Hamada, M., Murase, N., Hasan, A., Balaratnam, M., & Rothwell, J. C. (2013). The role of interneuron networks in driving human motor cortical plasticity. Cerebral Cortex, 23(7), 1593–1605. doi: 10.1093/cercor/bhs147

Huang, Y. Z., Edwards, M. J., Rounis, E., Bhatia, K. P., & Rothwell, J. C. (2005). Theta burst stimulation of the human motor cortex. Neuron, 45(2), 201–206. doi: 10.1016/j.neuron.2004.12.033

Jeffreys, H. (1939/1961). The theory of probability (1st/3rd Edn. ed.). Oxford, England: Oxford University Press.

Jenmalm, P., Schmitz, C., Forssberg, H., & Ehrsson, H. H. (2006). Lighter or heavier than predicted: neural correlates of corrective mechanisms during erroneously programmed lifts. Journal of Neuroscience, 26(35), 9015–9021. doi: 10.1523/JNEUROSCI.5045-05.2006

Johansson, R. S., & Westling, G. (1988). Coordinated isometric muscle commands adequately and erroneously programmed for the weight during lifting task with precision grip. Experimental Brain Research, 71(1), 59–71.

Kourtzi, Z., & Kanwisher, N. (2001). Representation of perceived object shape by the human lateral occipital complex. Science, 293(5534), 1506–1509. doi: 10.1126/science.1061133

Li, Y., Randerath, J., Goldenberg, G., & Hermsdorfer, J. (2007). Grip forces isolated from knowledge about object properties following a left parietal lesion. Neuroscience Letters, 426(3), 187–191. doi: 10.1016/j.neulet.2007.09.008

Li, Y., Randerath, J., Goldenberg, G., & Hermsdorfer, J. (2011). Size-weight illusion and anticipatory grip force scaling following unilateral cortical brain lesion. Neuropsychologia, 49(5), 914–923. doi: 10.1016/j.neuropsychologia.2011.02.018

Lopez-Alonso, V., Cheeran, B., Rio-Rodriguez, D., & Fernandez-Del-Olmo, M. (2014). Inter-individual variability in response to non-invasive brain stimulation paradigms. Brain Stimul, 7(3), 372–380. doi: 10.1016/j.brs.2014.02.004

Milner, A. D., & Goodale, M. A. (2008). Two visual systems re-viewed. Neuropsychologia, 46(3), 774–785. doi: 10.1016/j.neuropsychologia.2007.10.005

Monaco, S., Chen, Y., Medendorp, W. P., Crawford, J. D., Fiehler, K., & Henriques, D. Y. (2014). Functional magnetic resonance imaging adaptation reveals the cortical networks for processing grasp-relevant object properties. Cerebral Cortex, 24(6), 1540–1554. doi: 10.1093/cercor/bht006

Monaco, S., Sedda, A., Cavina-Pratesi, C., & Culham, J. C. (2015). Neural correlates of object size and object location during grasping actions. European Journal of Neuroscience, 41(4), 454–465. doi: 10.1111/ejn.12786

Murata, A., Gallese, V., Luppino, G., Kaseda, M., & Sakata, H. (2000). Selectivity for the shape, size, and orientation of objects for grasping in neurons of monkey parietal area AIP. Journal of Neurophysiology, 83(5), 2580–2601. doi: 10.1152/jn.2000.83.5.2580

Nowak, D. A., Voss, M., Huang, Y. Z., Wolpert, D. M., & Rothwell, J. C. (2005). High-frequency repetitive transcranial magnetic stimulation over the hand area of the primary motor cortex disturbs predictive grip force scaling. European Journal of Neuroscience, 22(9), 2392–2396. doi: 10.1111/j.1460-9568.2005.04425.x

Oldfield, R. C. (1971). The assessment and analysis of handedness: the Edinburgh inventory. Neuropsychologia, 9(1), 97–113.

Plaisier, M. A., & Smeets, J. B. (2015). Object size can influence perceived weight independent of visual estimates of the volume of material. Sci Rep, 5(February), 17719. doi: 10.1038/srep17719

Rice, N. J., Tunik, E., & Grafton, S. T. (2006). The anterior intraparietal sulcus mediates grasp execution, independent of requirement to update: new insights from transcranial magnetic stimulation. Journal of Neuroscience, 26(31), 8176–8182. doi: 10.1523/JNEUROSCI.1641-06.2006

Rossi, S., Hallett, M., Rossini, P. M., Pascual-Leone, A., & Group, T. S. o. T. C. (2009). Safety, ethical considerations, and application guidelines for the use of transcranial magnetic stimulation in clinical practice and research. Clin Neurophysiol, 120(12), 2008–2039. doi: 10.1016/j.clinph.2009.08.016

Saccone, E. J., & Chouinard, P. A. (2019). The influence of size in weight illusions is unique relative to other object features. Psychon Bull Rev, 26(1), 77–89. doi: 10.3758/s13423-018-1519-5

Schmitz, C., Jenmalm, P., Ehrsson, H. H., & Forssberg, H. (2005). Brain activity during predictable and unpredictable weight changes when lifting objects. Journal of Neurophysiology, 93(3), 1498–1509. doi: 10.1152/jn.00230.2004

Trewartha, K. M., & Flanagan, J. R. (2017). Linking actions and objects: Context-specific learning of novel weight priors. Cognition, 163, 121–127. doi: 10.1016/j.cognition.2017.02.014

Tunik, E., Frey, S. H., & Grafton, S. T. (2005). Virtual lesions of the anterior intraparietal area disrupt goal-dependent on-line adjustments of grasp. Nature Neuroscience, 8(4), 505–511. doi: 10.1038/nn1430

van Nuenen, B. F. L., Kuhtz-Buschbeck, J., Schulz, C., Bloem, B. R., & Siebner, H. R. (2012). Weight-specific anticipatory coding of grip force in human dorsal premotor cortex. Journal of Neuroscience, 32(15), 5272–5283. doi: 10.1523/JNEUROSCI.5673-11.2012

van Polanen, V., & Davare, M. (2015a). Interactions between dorsal and ventral streams for controlling skilled grasp. Neuropsychologia, 79(Pt B), 186–191. doi: 10.1016/j.neuropsychologia.2015.07.010

van Polanen, V., & Davare, M. (2015b). Sensorimotor Memory Biases Weight Perception During Object Lifting. Front Hum Neurosci, 9, 700. doi: 10.3389/fnhum.2015.00700

